# Pial collaterals propagate neurovascular responses across functional units

**DOI:** 10.64898/2025.12.09.692989

**Authors:** Chaim Glück, Matthias T. Wyss, Alba Vieites-Prado, Jacqueline Condrau, Jeanne Droux, Quanyu Zhou, Zhenyue Chen, Nicolas Renier, Daniel Razansky, Aiman Saab, Susanne Wegener, Bruno Weber, Mohamad El Amki

**Author notes:** These authors contributed equally to this work (Bruno Weber, Mohamad El Amki).

## Abstract

Neurovascular coupling refers to the regulation of cerebral blood flow in response to neuronal activity, a process forming the basis for multiple functional neuroimaging techniques. However, the mechanisms that coordinate neurovascular responses across distinct cortical arterial territories remain poorly understood. Here we show that pial collaterals, arterio-arterial connections traditionally considered dormant under physiological conditions, serve as active conduits that propagate neurovascular signals between adjacent cortical territories and functional units. We demonstrate that neuronal activity in the somatosensory cortex triggers direct vascular signaling via endothelial Kir2.1 channels through pial collaterals, synchronizing blood flow responses beyond the activated area. The loss of pial collaterals abolished inter-territorial coupling without affecting local neurovascular responses. These results reveal that pial collaterals are critical regulators of cerebral perfusion.

## Main Text

Neurovascular coupling ensures that cerebral blood flow is finely matched to the metabolic demands of active neurons ^1–5^. Understanding the specific responses of brain vascular networks to neuronal activity is the basis of functional magnetic resonance imaging (fMRI) and remains a fundamental challenge in neurobiology ^5,6^. During neuronal activity, neurons generate localized changes in the extracellular environment, leading to increased blood flow at the capillary level, which propagates upstream in arterioles ^3,7–10^. While capillary-to-arteriole signaling is central to this process, current models assume that hyperemic responses remain confined within discrete arterial territories ^9,11^. Whether neuronal activity can initiate vascular responses across territorial boundaries remains unknown.

Pial collaterals, which connect distal branches of adjacent arterial trees, provide anatomical routes for inter-territorial flow ^12–15^. Although crucial during stroke for collateral perfusion, these vessels are considered dormant under physiological conditions ^16–18^. It is unknown whether they contribute to the propagation of neurovascular responses. The amount of connectivity in the arterial connectome is defined by the number of pial collaterals, which differ in density between individual brains ^19^. This variability in vascular architecture raises intriguing questions about how connectivity maps in the arterial network impact the propagation of functional hyperemic responses. Do pial collaterals respond to increased neuronal activity? To what extent does the functional hyperemic response propagate through the pial arterial connectome? Once the vasodilatory response reaches the distal end of an arterial network, does it stop, or does it travel via collaterals to neighboring networks? Finally, are the pial collaterals routes for vascular signaling that enable inter-territorial neurovascular responses?

Here, we show that pial collaterals actively propagate vascular responses between adjacent arterial territories in the somatosensory cortex. This inter-territorial signaling is triggered by neuronal activity and requires endothelial Kir2.1 channels but is independent of pressure gradients or local neuronal activity spillover. These findings identify pial collaterals as active elements of the neurovascular signaling network, revealing a vascular mechanism for long-range coordination of cerebral blood flow across functional units.

## Results

### Pial collaterals are connectivity hubs providing inter-territorial connections within the arterial connectome

To map inter-territorial arterial connectivity, we applied tissue clearing and volumetric imaging with immunolabelling for smooth muscle actin (α-SMA), CD31, and Podocalyxin to visualize the cortical arterial connectome in 3D ^20^. The anterior, middle, and posterior cerebral arteries (ACA, MCA, and PCA, respectively) were readily identified, giving rise to distal branches that defined non-overlapping arterial territories (Fig. 1a). In watershed regions, pial collaterals formed anastomotic connections between terminal branches of different arteries, with the highest number observed between MCA and ACA territories. Each hemisphere contained a median of ten such connections, though numbers varied across individual brains (Fig. 1a). These inter-territorial links define the degree of connectivity within the arterial network.

**Fig. 1.**
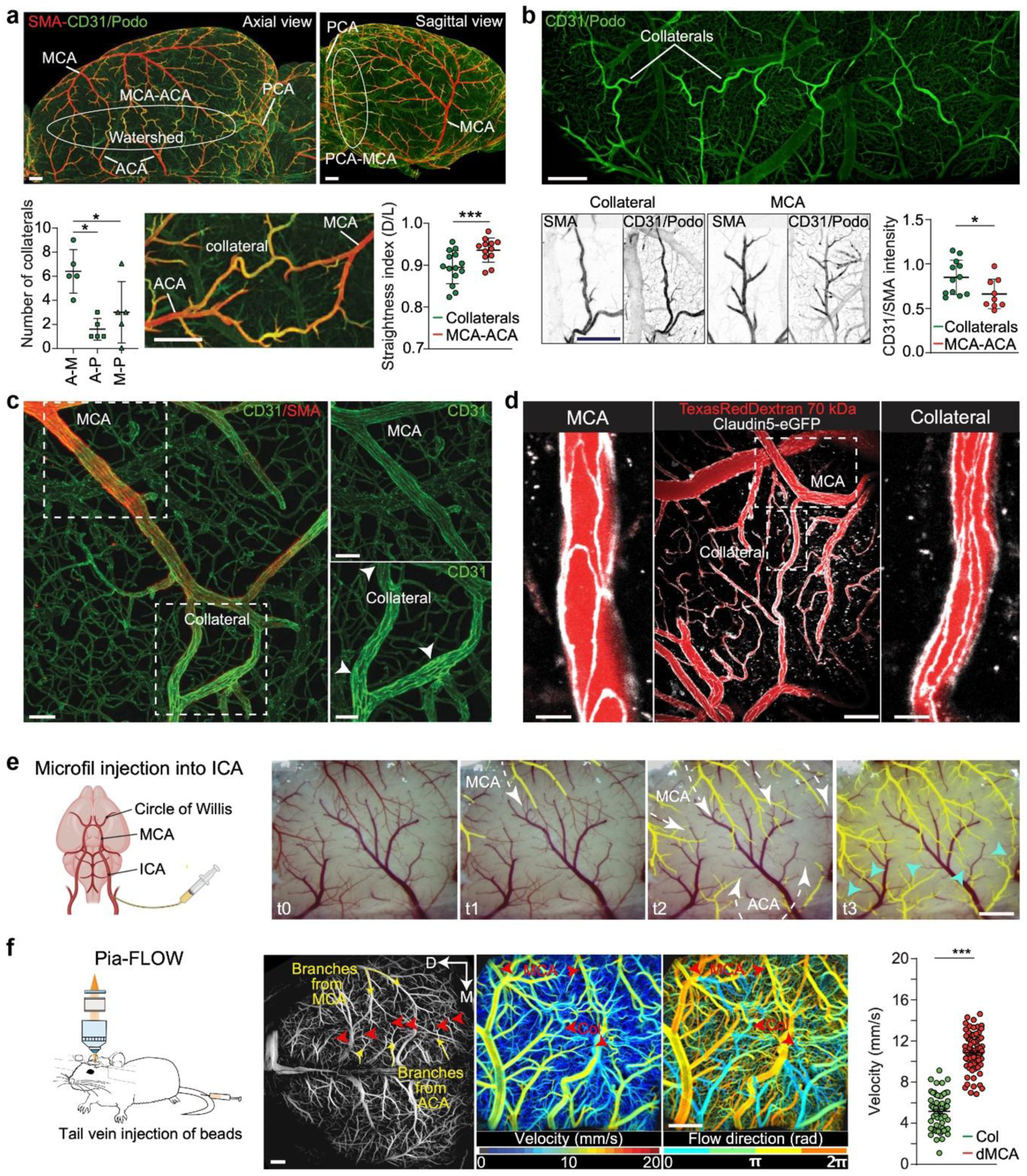
Pial collaterals are connectivity hubs providing inter-territorial connections within the arterial connectome. (**a**) (Top) Axial and sagittal views of a brain hemisphere processed with iDISCO+. Arterial walls are labeled for smooth muscle actin (α-SMA) and the vascular endothelium is visualized with a combination of CD31. The main pial arteries, MCA, ACA and PCA are interconnected by anastomosed collaterals in the watershed area; scale bars: 500°µm. (Bottom) Quantification of pial collaterals connecting the different vascular territories. A-M: ACA-MCA, A-P: ACA-PCA and M-P: MCA-PCA. Tortuosity of arterial collaterals. Pial artery segments are significantly straighter than collaterals (p = 0.032, Two-way mixed-effects ANOVA); scale bars: 500 µm. (**b**) (Top) Image of a CD31/Podocalyxin (Podo) staining, highlighting increased immunoreactivity in collaterals compared to MCA segments. (Bottom) SMA and CD31/Podo stainings comparing collateral with MCA segments. Anastomosed collaterals exhibited enriched expression of endothelial markers (p=0.013, Two-way mixed-effects ANOVA); scale bars: 300 µm. (**c**) High magnification light sheet microscopy images of a CD31 and SMA staining in collaterals compared to MCA segments, scale bars: 50 µm. (**d**) *In vivo* two-photon image of the cortical vasculature of a Claudin5-eGFP mouse, showing differences in endothelial cell shape in collaterals (right panel) compared to MCA segments (left panel). Blood plasma is stained with TexasRed Dextran 70 kDa and tight-junctions (Claudin5) are labelled with eGFP, scale bars: 50 µm (left and right panel), 100 µm (middle panel). (**e**) Injection of Microfil into the internal carotid artery (ICA) to cast the arterial network. Images show timeseries (t0-t3) with gradual filling of the arterial network and respective flow-directions (indicated by white dashed arrows), t1=2s, t2=4s and t3=8s. Cyan arrowheads indicate the position of collaterals; scale bar: 500 µm. (**f**) (Left) Scheme of the experimental Pia-FLOW setup. (Middle) Image of whole pial vasculature with corresponding velocity and angle maps. Red arrowheads indicate collaterals and red arrowheads indicate MCA and ACA. (Right) Quantification of blood flow velocities comparing Collaterals (Col) and distal MCA (dMCA) branches; scale bars: 500 µm. (A-E) N = 3-5 mice.

Collateral segments displayed increased tortuosity relative to non-anastomosed distal arterioles (Fig. 1a). Analysis of cellular composition revealed no significant difference in α-SMA signal, but a marked enrichment of endothelial markers in collateral segments compared to adjacent arterioles. As a result, the endothelial-to-smooth muscle signal ratio was significantly higher in anastomotic segments (Fig. 1b-d), consistent with distinct endothelial specialization that may support enhanced sensitivity to hemodynamic forces. We hypothesize that this enables bidirectional flow through collaterals, allowing them to dynamically adjust to the metabolic and perfusion needs of different cortical regions.

Collateral vessels also exhibited unique flow dynamics. Microfil injections into the carotid artery revealed sharp boundaries between opposing ACA and MCA flow domains, with collaterals appearing less pressurized and sparsely filled, indicative of low flow under physiological conditions (Fig. 1e and Extended Data movie S1). In a complementary experiment, to assess collateral dynamics in real time and to get quantitative measures of flow in the entire arterial connectome, we used Pia-FLOW, an in vivo widefield fluorescence localization microscopy ^12,21^ (Fig. 1f). Flow velocity was significantly lower in pial collaterals than in terminal branches of the ACA or MCA (Fig. 1f), consistent with previous reports ^13,17,22^ and confirming their role as low-flow, high-resistance conduits under resting conditions.

### Collaterals mediate inter-territorial neurovascular coupling

To test whether pial collaterals are functionally active under physiological conditions, we performed in vivo two-photon imaging in awake Acta2-RCaMP1.07 reporter mice. In this line, smooth muscle cells express the red genetically encoded calcium indicator RCaMP1.07, enabling real-time visualization of intracellular Ca²⁺ dynamics. Under resting conditions, collaterals displayed irregular, intermittent, and turbulent blood flow (Fig. 2a-c and Extended Data movies S2,S3). Calcium activity in collateral smooth muscle cells was comparable to that observed in the distal pial arteriole (Fig. 2d, Extended Data Fig. 1), indicating that collaterals are sites of spontaneous vascular signaling and suggesting an overlooked physiological function.

**Fig. 2.**
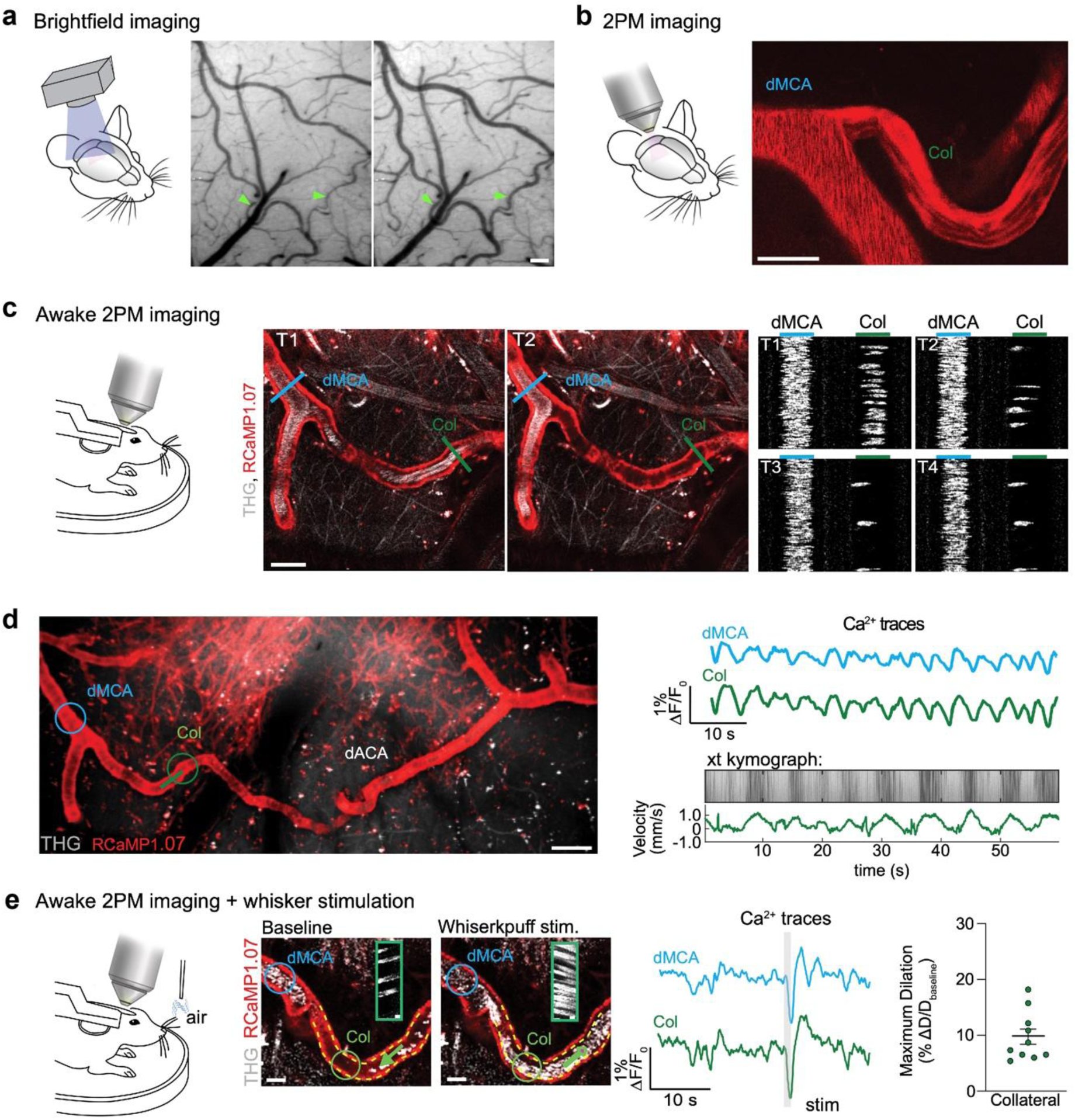
Collaterals mediate functional hyperemic responses between arterial networks. (**a**) Brightfield images through a cranial window showing different perfusion states of pial collaterals (indicated by green arrowheads); scale bar: 100 µm. (**b**) Two-photon (2PM) image of a distal MCA (dMCA) and collateral (col). Blood flow in the col shows an irregular blood flow pattern compared to blood flow in the dMCA branch. Blood plasma is stained with TexasRed Dextran 70 kDa; scale bar: 50 µm. (**c**) (Left) Scheme of the experimental setup. (Middle) Awake two-photon images of distal MCA - collateral branches of an Acta2-RCaMP1.07 mouse at different timepoints (T1, T2). Blue and green lines indicate the position of line scans. (Right) Line scan images at different timepoints (T1-T4); scale bars: 50 µm. (**d**) (Left) Awake two-photon images of the MCA-collateral-ACA arterial branch of an Acta2-RCaMP1.07 mouse. Circles and lines indicate positions of recordings for Ca^2+^ and blood flow. (Right) Ca^2+^ - traces of a dMCA and col (top) and kymograph (bottom) are shown. dACA = distal ACA, THG = third harmonic generation; scale bar: 50 µm. (**e**) (Left) Awake two-photon images of an MCA-collateral branch before and during whisker air-puff stimulation. Circles and arrows indicate positions of recordings for Ca^2+^ and blood flow. Ca^2+^ - traces (on top) and kymograph are shown. On the right, quantification of the fractional dilation of collaterals upon whisker air puff stimulation, yellow dashed lines indicate the baseline vessel lumen diameter; scale bar: 25 µm. N = 10 collaterals from three mice.

We next investigated whether collaterals actively contribute to neurovascular coupling. We focused on the barrel cortex, a sensory domain of the primary somatosensory cortex that receives input from the whisker pad and is exclusively supplied by the MCA arterial network ^23–26^. Whisker stimulation in awake Acta2-RCaMP1.07 mice triggered a robust decrease in intracellular Ca²⁺ in smooth muscle cells of arterioles within the barrel cortex, followed by vasodilation measured through red blood cell displacement (Fig. 2e and movie S4). Notably, similar RCaMP1.07 and vasodilatory responses were also detected in collateral vessels connected to the stimulated MCA branches, with no significant differences in response amplitude or timing. These findings indicate that collaterals are recruited during neuronal activity and function as integral components of the neurovascular response.

To assess the extent of vascular response propagation in space and across large distances, we employed widefield optical imaging. We first defined the neuronal responses and mapped the responsive whisker region by voltage sensitive dye (VSD) imaging which provides a fast and precise visualization of the ensemble membrane potential dynamics (Fig. 3a, Extended Data Fig. 2). Correspondingly, neuronal calcium activity in Thy1-GCaMP6f mice confirmed localized activation within the MCA-supplied territory (Fig. 3a), consistent with anatomical mapping. Overlaying vascular topology maps onto the activated region confirmed that whisker input engaged networks exclusively supplied by the MCA, in accordance with previous reports^26^.

**Fig. 3.**
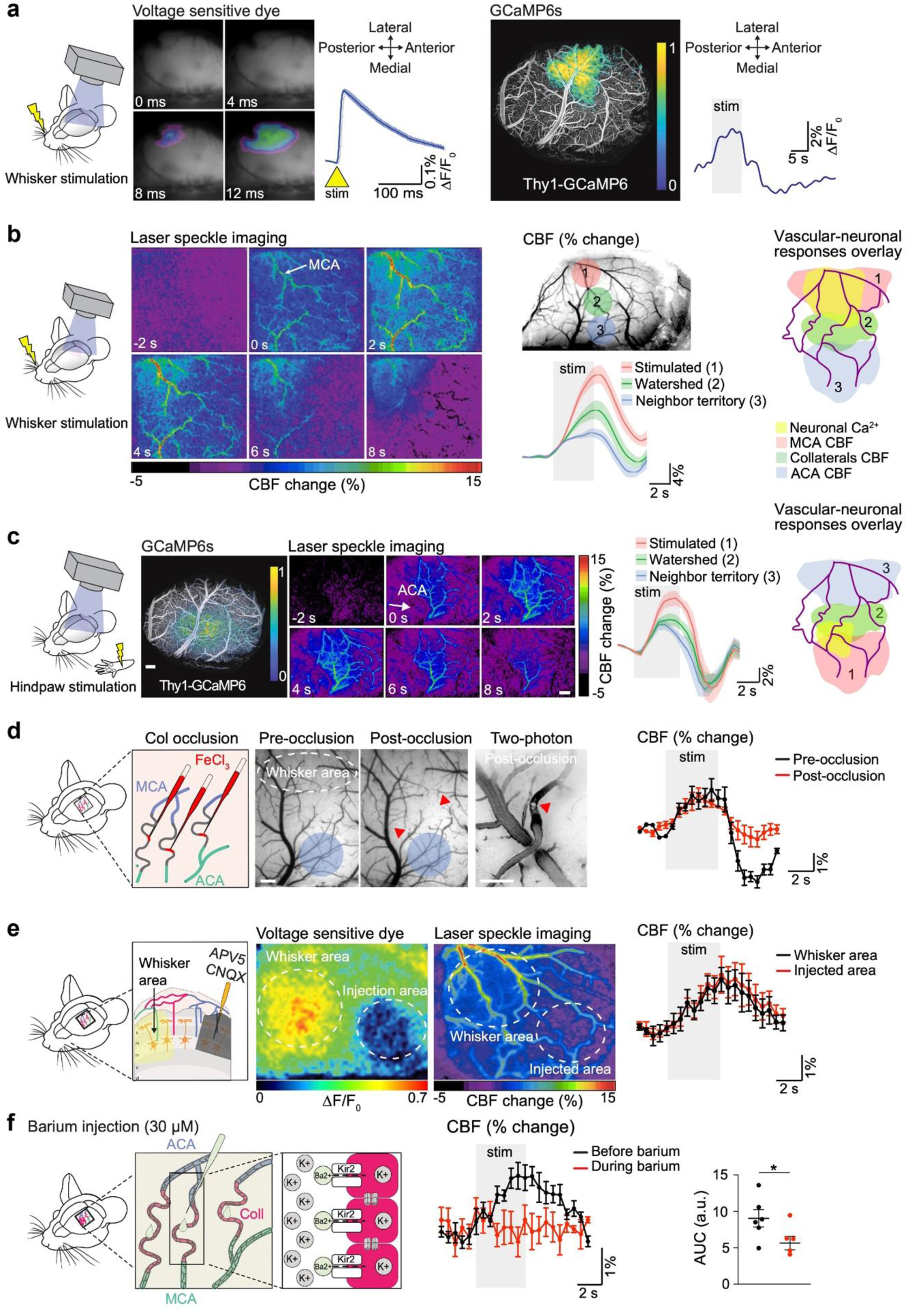
Collateral transmission of neurovascular coupling is mediated by ECs signaling. (a) Mapping whisker area by measuring the whisker stimulation response of a C57BL/6 mouse with voltage-sensitive dye (VSD) imaging (left) and GCaMP6s fluorescence imaging (right). (b) Laser speckle imaging (LSI) showing cerebral blood flow (CBF) changes upon whisker stimulation. Positions of region of interest (ROI) and CBF responses are shown (N = 3 mice). On the right, scheme of the arterial territories and stimulation area (yellow); scale bar: 500 µm. (c) Mapping of hindpaw area by GCaMP6s fluorescence imaging and LSI of CBF response upon hindpaw stimulation. On the right, scheme of the arterial territories and stimulation area (yellow); scale bars: 500 µm. (**d**) Occlusion of collaterals by FeCl_3_ (indicated by red arrowheads) and measurement of CBF response in neighboring territory. On the right CBF response curve in the neighbor territory upon whisker stimulation. N = 4 mice; scale bars: 100 µm. (**e**) Blockade of neuronal firing with APV/CNQX and measurement of CBF response in the injection area. (Right) CBF response curve in the injected area upon whisker stimulation. N = 5 mice; scale bars: 500 µm. (**f**) Blockade of endothelial Kir2 channels by topical Barium (BaCl_2_; 30 µM) application and measurement of CBF response in the neighboring area. On the right CBF response curve in the adjacent area upon whisker stimulation. N = 6 mice.

Vascular imaging revealed that sensory-evoked hyperemia initiated within the MCA territory propagated across collateral vessels into the neighboring ACA domain (Fig. 3b). Although delayed and smaller in amplitude, the hemodynamic response in ACA branches indicated that functional signals can cross inter-territorial boundaries via collaterals, even between vascular domains supplied by opposing flow directions.

To test whether this propagation is bidirectional, we stimulated the hindlimb, which is served by the ACA arterial network. Hindlimb stimulation evoked local neurovascular responses in the hindlimb cortex and, notably, also increased blood flow in the adjacent MCA domain (Fig. 3c). The amplitude of the response was highest in ACA branches but also present in MCA branches connected via collaterals, indicating signal transmission in the reverse direction. Despite opposing baseline flow directions between ACA and MCA territories, the functional hyperemic response propagated across collaterals in both directions with no orientation selectivity. These data demonstrate that pial collaterals serve as conduits for bidirectional propagation of neurovascular signals between distinct arterial territories

### Endothelial signaling mediates collateral-driven vascular responses

To determine whether blood flow itself drives neurovascular signal propagation through collaterals, we selectively occluded MCA-ACA collateral segments around the hindlimb cortex. We established a targeted clot induction protocol by applying ferric chloride (FeCl₃, 10%) to the cortical surface using a glass micropipette. Contact with the pial surface near identified collaterals induced localized thrombosis, as previously described ^27,28^. Two-photon imaging confirmed intraluminal clot formation and complete cessation of flow in the occluded vessels (Fig. 3d).

To assess whether collateral flow was required for signal transmission, we performed laser speckle imaging (LSI) during whisker stimulation. We monitored hemodynamic responses in both the barrel cortex and the neighboring ACA territory. Despite successful collateral occlusion, delayed hemodynamic responses persisted in the ACA region. These findings argue against passive regulation of blood flow and suggest that vascular responses in adjacent territories can occur independently of collateral perfusion.

To rule out the possibility that neural activity was spreading to the ACA territory, we selectively suppressed synaptic transmission in the hindlimb cortex. A mixture of CNQX and AP5, antagonists of AMPA and NMDA receptors respectively, was injected locally to block glutamatergic signaling in the non-stimulated region (Fig. 3e) ^29^. This intervention left the barrel cortex intact while silencing potential neuronal co-activation in the ACA domain. VSD imaging confirmed the absence of stimulus-evoked membrane depolarization in the drug-injected area. Nevertheless, hemodynamic responses were still observed in the ACA territory during whisker stimulation, indicating that vascular signal propagation was not a secondary effect of neuronal spread.

We next tested whether signaling within the endothelium mediates this long-range vascular coordination. Prior studies implicated Kir2.1 potassium channels in endothelial cell (EC)-mediated retrograde signaling during neurovascular coupling ^30–32^. To test this, we applied the Kir2.1 blocker barium chloride (BaCl₂, 30 µM) topically to the exposed pial surface over the MCA-ACA border. Inhibition of Kir2.1 channels abolished hemodynamic responses in neighboring territories during sensory stimulation (Fig. 3f). These findings support a model in which EC-based signaling, rather than flow or neuronal spread, underlies collateral-mediated transmission of neurovascular responses across arterial networks.

### Collateral deficiency restricts inter-territorial vascular signaling

To assess whether arterial connectivity is required for long-range neurovascular responses, we examined Rabep2-/- mice, which lack pial collaterals due to a developmental defect in collateral formation ^13,33^. Using Pia-FLOW and two photon imaging, we confirmed a marked reduction in collateral number, particularly between the MCA and ACA territories (Fig. 4a). To verify that local neurovascular coupling remained intact, we first examined sensory-evoked neuronal activity in the somatosensory cortex using VSD imaging. Rabep2-/- and control mice exhibited comparable neuronal response amplitude and kinetics (Extended Data Fig. 3). Following whisker stimulation, blood flow increased within the MCA-supplied barrel cortex similarly in both genotypes, as measured by LSI (Fig. 4b-c), indicating preserved local vascular responses.

**Fig. 4.**
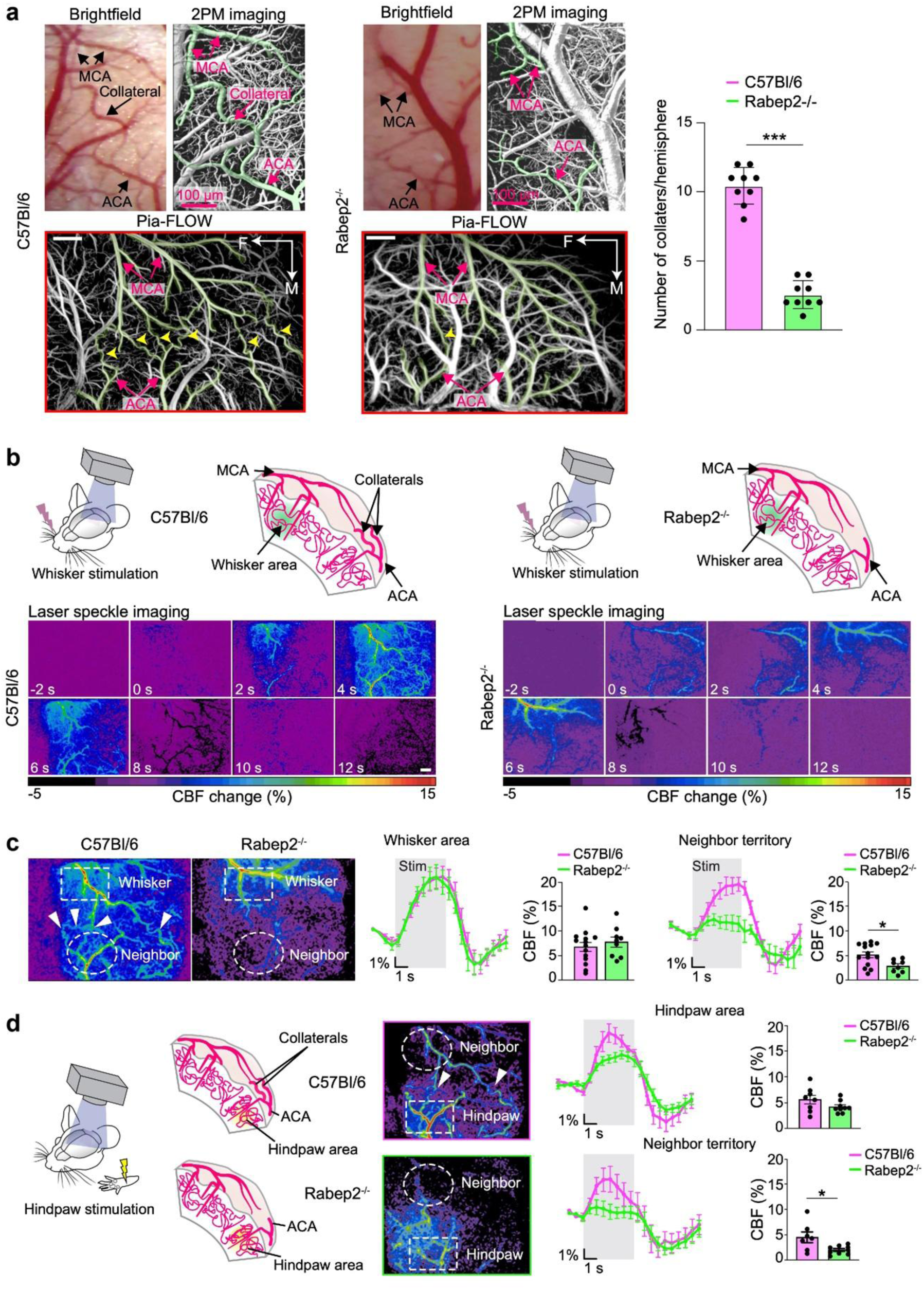
Genetic depletion of pial collaterals impairs the neurovascular response. (**a**) Vascular images (brightfield, two-photon and Pia-FLOW) comparing pial vasculature of C57BL/6 with rabep2^-/-^ mice. On the right quantification of the number of collaterals in respective mouse strains. N = 9 mice per strain. Scale bars: 100 µm. (**b**) (Left) Schematic diagrams illustrating the brain region of interest as well as the vasculature in the C57BL/6 mice (top). LSI images of C57BL/6 mice upon whisker stimulation (Bottom). (Right) Schematic diagrams illustrating the brain region of interest as well as the vasculature in the rabep2^-/-^ mice (top). LSI images of rabep2^-/-^ mice upon whisker stimulation (Bottom). (**c**) Quantification of CBF change upon whisker stimulation in the stimulated and neighboring areas comparing C57BL/6 with rabep2^-/-^ mice. N = 9-14 mice per group. (**d**) (Left) Schematic diagrams illustrating the brain region of interest as well as the vasculature in the C57BL/6 mice (top) and rabep2^-/-^ mice (Bottom). (Middle) LSI images of C57BL/6 with rabep2^-/-^ mice upon hindpaw stimulation. (Right) Quantification of CBF change upon hindpaw stimulation in stimulated and neighboring areas. N = 7-9 mice per group.

We then tested whether hemodynamic signals spread across arterial boundaries. In control mice, whisker and hindlimb stimulation produced vascular responses that extended beyond the initiating territory, consistent with collateral-mediated propagation. In contrast, Rabep2-/- mice showed an 84% reduction in hemodynamic spread after whisker stimulation and a 94% reduction after hindlimb stimulation (Fig. 4c). These deficits resembled the effects of pharmacological blockade of endothelial signaling (Fig. 3f), suggesting that collateral absence disrupts long-range vascular coordination.

Importantly, resting-state flow dynamics and stimulus-evoked arteriolar dilation within the activated territory were similar between Rabep2-/- and control animals, as confirmed by two-photon microscopy and LSI (Extended Data Fig. 3). Thus, the diminished inter-territorial response is not due to global vascular impairments, but rather to loss of anatomical connectivity. Together, these findings demonstrate that pial collaterals are essential for the propagation of neurovascular signals across the arterial network.

## Discussion

We identified a previously unrecognized physiological role for pial collaterals in enabling long-range vascular communication via EC signaling. Far from being passive conduits, these vessels actively propagate functional hyperemia across distinct arterial networks, forming a dynamic part of the neurovascular connectome. By coupling neuronal activity to widespread coordinated vascular responses, collaterals integrate the local and global regulation of cerebral blood flow. Although pial collaterals have been traditionally viewed as dormant under normal conditions and only recruited in disease states such as stroke ^18,34^ or Moyamoya disease ^13,14,34–36^, our results redefine them as active participants in vascular signaling during physiological brain function. We showed that neuronal activation initiates vasodilation in the local pial network and that this response is transmitted to adjacent arterial territories via collaterals. This process is not driven by passive blood redistribution or hemodynamics, but rather depends on EC signaling, as shown by the abolition of inter-territorial responses upon Kir2.1 channel blockade. Mechanistically, these findings expand the established role of ECs in neurovascular coupling, previously demonstrated primarily at the capillary level ^30–32^. Our work is in line with recent work showing that ECs in arteries are functionally coupled and serve as a signaling highway to enable rapid vasodilation propagation during neurovascular coupling ^37^. Moreover, our data support a model in which ECs in superficial pial arteries act as electrical conduits, transmitting vasodilatory signals across arterial domains through collaterals. This form of vascular crosstalk enables synchronized hemodynamic responses beyond the boundaries of individual arterial territories. Notably, these signals persisted even when neighboring neuronal activity was pharmacologically silenced, further supporting a vascular origin of the response. Our findings support a recent study showing that functional hyperemic responses may be highly heterogeneous across individual vessel segments and do not necessarily reflect local neuronal activity ^38^.

Experiments in Rabep2⁻/⁻ mice, which lack pial collaterals due to developmental defects ^13,33^, provided further support. These animals exhibited normal initiation of neurovascular responses in the stimulated cortex but lacked the propagation of vascular responses to neighboring territories. Baseline perfusion and capillary flow remained unaffected, indicating that collaterals are dispensable for resting blood flow but are required for the spatial extension of functional hyperemia.

These findings carry broader implications for interpreting neuroimaging signals and understanding individual variability in neurovascular physiology. Variations in arterial connectivity, collateral number and architecture may underlie differences in BOLD-fMRI responses, especially in contexts where the spatial extent of blood flow changes is used to infer patterns of neural activity ^9,39^. Moreover, our data challenge the classical view that neurovascular coupling merely reflects immediate local metabolic demands. Instead, they suggest that pial collaterals provide a structural basis for redistributing blood flow beyond locally active regions, potentially tuning perfusion according to network-level needs.

Finally, this study opens new avenues for research into how vascular architecture contributes to brain resilience. Future studies should explore how aging, genetic variation, or disease conditions affect the signaling capacity and plasticity of collaterals, and whether similar inter-territorial signaling mechanisms operate in other vascular beds beyond the cortex.

## Material and methods

### Animals

For all experiments, C57BL/6 (Charles Rivers, no. 000664), Acta2-RCaMP1.07 (Jackson, No. 028345), and Rabep2-/- mice, mixed female/male, between 3 to 4 months of age were used. There was no influence/association of sex on the findings. The animals had free access to water and food and an inverted 12-hour light/dark cycle to perform experiments during the active phase. All experiments were approved by the local veterinary authorities in Zurich and conformed to the guidelines of the Swiss Animal Protection Law, Veterinary Office, Canton of Zurich (Act of Animal Protection 16 December 2005 and Animal Protection Ordinance 23 April 2008), animal welfare assurance numbers ZH165/19, ZH224/15, and ZH152/2021.

### Samples staining and iDISCO+ clearing

Whole brain vasculature staining was performed using the iDISCO+ protocol previously described ^20,40^ with minimal modifications. All the steps of the protocol were done at room temperature with gentle shaking unless otherwise specified. All the buffers were supplemented with 0.01% sodium azide (Sigma-Aldrich, Germany) to prevent bacterial and fungi growth. Perfused brains were dehydrated in an increasing series of methanol (Sigma-Aldrich, France) dilutions in water (washes of 1 hour in methanol 20%, 40%, 60%, 80% and 100%). An additional wash of 2 hours in methanol 100% was done to remove residual water. Once dehydrated, samples were incubated overnight in a solution containing 66% dichloromethane (Sigma-Aldrich, Germany) in methanol, and then washed twice in methanol 100% (4 hours each wash). Samples were then bleached overnight at 4°C in methanol containing 5% hydrogen peroxide (Sigma-Aldrich). Rehydration was performed by incubating the samples in methanol at 60%, 40%, and 20% concentrations (each for 1 hour). After methanol pretreatment, samples were washed in PBS twice for 15 minutes, 1 hour in PBS containing a 0,2% of Triton X-100 (Sigma-Aldrich), and further permeabilized by a 24 hours incubation at 37°C in Permeabilization Solution, composed by 20% dimethyl sulfoxide (Sigma-Aldrich), 2,3% Glycine (Sigma-Aldrich, USA) in PBS-T. To initiate the immunostaining process, samples were first blocked with 0.2% gelatin (Sigma-Aldrich) in PBS-T for 24 hours at 37°C. The same blocking buffer was used to prepare antibody solutions. A combination of primary antibodies targeting different components of the vessel’s walls was used to achieve continuous immunostaining. Alpha Smooth Muscle Actin (αSMA or Acta2) antibody was used to label the artery’s wall, and antibodies to Podocalyxin and CD31 were combined to stain the full capillary net and large veins. Primary antibodies were incubated for 10 days at 37°C with gentle shaking, then washed in PBS-T (twice for 1 hour and then overnight), and finally incubated for 10 days with secondary antibodies. Secondary antibodies conjugated to Alexa 647 were used to detect Podocalyxin and CD31, while arteries were stained with secondary antibodies conjugated to Alexa 555. After immunostaining, the samples were washed in PBS-T (twice 1 hour and then overnight), dehydrated in a methanol/water increasing concentration series (20%, 40%, 60%, 80%, 100% one hour each followed by overnight incubation of methanol 100%), followed by a wash in 66% dichloromethane – 33% methanol for 3 hours. Methanol was washed out with two final washes in dichloromethane 100% (15 min each), and finally, the samples were cleared and stored in dibenzyl ether (Sigma-Aldrich) until light sheet imaging.

### Anaesthesia

For head-post and cranial window implantation, mice were anesthetized intraperitoneally with a mixture of fentanyl (0.05 mg/kg bodyweight; Sintenyl, Sintetica), midazolam (5 mg/kg bodyweight; Dormicum, Roche), and medetomidine (0.5 mg/kg bodyweight; Domitor, Orion Pharma). Throughout anaesthesia, a facemask provided oxygen at a rate of 500 ml/min. For two-photon imaging under anaesthesia, animals were induced with isoflurane 4%, maintained at 1.2% with continuous supply of an oxygen/air mixture (30%/70%). For LSI recordings, anaesthesia was changed to a mixture of fentanyl (0.05 mg/kg bodyweight; Sintenyl, Sintetica), midazolam (5 mg/kg bodyweight; Dormicum, Roche), and medetomidine (0.5 mg/kg bodyweight; Domitor, Orion Pharma). The core temperature of the animals was maintained at 37°C using a homeothermic blanket heating system (Harvard Apparatus) throughout all surgical and experimental procedures. The animaĺs head was fixed in a stereotaxic apparatus, and the eyes were kept moist with a vitamin A eye cream (Bausch & Lomb).

### Head-Post Implantation

A bonding agent (Gluma Comfort Bond; Heraeus Kulzer) was applied to the cleaned skull and polymerized with a handheld blue light source (600 mW/cm2; Demetron LC). A custom-made aluminum head post was connected to the bonding agent with dental cement (EvoFlow; Ivoclar Vivadent AG) for stable and reproducible fixation in the microscope setup. The skin lesion was treated with an antibiotic cream (Fucidin®, LEO Pharma GmbH) and closed with acrylic glue (Histoacryl, B. Braun). After surgery, animals were kept warm and received analgesics (buprenorphine 0.1 mg/kg bodyweight; Sintetica).

### Cranial window surgery

A 4 × 4 mm craniotomy was performed above the somatosensory cortex (centered above the left somatosensory cortex) with a dental drill (Bien-Air). A square coverslip (3 × 3 mm, UQG Optics) was placed on the exposed dura mater and fixed to the skull with dental cement ^41^.

### Laser Speckle Imaging (LSI)

Cortical relative cerebral blood flow (CBF) changes were monitored before and during stimulation using a laser speckle contrast imaging monitor (FLPI, Moor Instruments, UK) ^42^. The acquisition was performed with a frame rate of 25 Hz. Afterwards, the LSCI images were exported and generated with arbitrary units in a 256-colour palette by the MOOR-FLPI software.

### Two-photon imaging

After cranial window implantation, mice recovered for at least 2 weeks prior to two-photon imaging. Imaging was performed using a custom-built two-photon laser scanning microscope (2PLSM) with a tunable pulsed laser (Chameleon Discovery TPC, Coherent Inc.) equipped with a 25× (W-Plan-Apochromat 25×/1.0 NA, Olympus) water immersion objective. During measurements, the animals were head-fixed and kept under anaesthesia as described above. Vasculature was labelled with intravenous injection of FITC–dextran (2-MDa, Sigma-Aldrich; FD2000S) or Texas red Dextran (70 kDa, Life Technologies catalogue number D-1864), ten minutes before imaging. Emission was detected using GaAsP photomultiplier modules (Hamamatsu Photonics) equipped with 475/64, 520/50, and 607/70 nm bandpass filters, and separated by 506, 560, and 652 nm dichroic mirrors (BrightLine; Semrock). The microscope was controlled by a customized version of ScanImage (r3.8.1; Janelia Research Campus 62).

For the acquisition of red blood cell (RBC) velocity and vessel diameters, line scans were performed in arterioles for 12.7 s. Three regions of interest (ROIs) were identified on the terminal distal branches of arterioles from the MCA. Three other ROIs were placed on collaterals, identified (1) as the arterio-arterial anastomosis connecting MCA to ACA and (2) by their typical tortuosity and low flow. The same vessels were evaluated before and after stimulation throughout the experiment.

Line scans were processed with a custom-designed image processing toolbox in MATLAB (https://github.com/EIN-lab/CHIPS). Vessel diameters were determined at full-width-half-maximum (FWHM) from a Gaussian fitted intensity profile drawn perpendicular to the vessel axis. RBC velocity flow was calculated with the Radon algorithm.

### Voltage-Sensitive Dye Imaging

The cortex was surgically exposed by bilateral craniotomy and durotomy, and the voltage-sensitive dye RH 1691 was applied for approximately 60 minutes, followed by wash-off with Ringer’s solution for approximately 30 minutes ^43,44^. VSD signal was imaged at 500 Hz using a MiCAM ULTIMA CMOS camera (SciMedia) coupled to a tandem lens array macroscope (50 mm f/1.4; Canon) under 630 nm excitation illumination. Sensory stimuli were delivered as for IOI. Data was subsequently processed in PMOD (PMOD Technologies). Response latency for each brain hemisphere was calculated as the time difference between stimulus onset and half-maximal activation. Similarly, interhemispheric latency was calculated as the time difference between half-maximal activation of each hemisphere.

### Intrinsic Optical Imaging

Cortical images were acquired using two 12-bit CCD cameras (pco.pixelfly VGA, PCO Imaging, Kelheim, Germany) attached to a motorized epifluorescence stereomicroscope (Leica MZ16 FA, Leica Microsystems, Heerbrugg, Switzerland) focused 0.5 mm below the cortical surface. Two-dimensional optical spectroscopy was performed using the method described by Dunn et al (2003) ^45–47^. The six wavelengths (560, 570, 580, 590, 600, and 610 nm, 10 nm FWHM) were produced with a monochromator (Polychrome V, Till Photonics, Grafelfing, Germany) and coupled in the microscope using an optical fiber. Images were acquired at 30 Hz, and the monochromator was synchronized with the image acquisition (each frame was acquired with a different illumination wavelength). The second camera was used to simultaneously measure CBF using dynamic laser speckle imaging. The method is described in detail elsewhere ^48^. A 785-nm laser light (TuiOptics, Munich, Germany) was shone onto the cortex and images were acquired at 30 Hz with an exposure time of 10 milliseconds.

### Pia-FLOW imaging and image reconstruction

Pia-FLOW imaging was performed as described previously ^12,21,49,50^. In brief, 50 μL of fluorescent beads (1 to 5 µm, FMOY, Cospheric, USA) in saline were injected via tail vein at a concentration of 2 × 10^8^ beads/mL. Epi-illumination of the sample was achieved with a 473 nm continuous wave (CW) laser (FPYL-473-1000-LED, Frankfurt Laser Company, Germany) beam that was expanded with a convex lens (LA4306, Thorlabs, USA) and an objective (CLS-SL, EFL = 70 mm, Thorlabs, USA). Backscattered fluorescence was collected with the same objective, focused with a tube lens (AF micro-Nikkor 105 mm, Nikon, Japan), and detected by a high-speed camera (pco.dimax S1, PCO AG, Germany). A longpass filter (FGL515, Thorlabs, USA) between the tube lens and camera filtered out the reflected excitation light. The system’s magnification was reduced to 1 for transcranial measurements to achieve an enlarged FOV by using a different camera lens (AF micro-Nikkor 60 mm, Nikon, Japan).

To generate density, flow velocity, and direction maps at high spatial resolution, the recorded widefield image stack was first denoised by subtracting a moving background calculated from the mean intensity of adjacent images to remove the static background stemming from the autofluorescence and dark noise of the camera. Subpixel localization was performed by extracting the centroid of bright spots detected using an adaptive threshold. Subsequently, the flowing fluorescent beads were tracked by identifying the same particle in different frames with the simpletracker algorithm (simpletracker.m. GitHub https://github.com/tinevez/simpletracker, 2019), with no gap filling and maximum linking distance corresponding to a velocity of 20 mm/s. Functional parameters, such as flow velocity and direction, were calculated with the given frame rate and relative displacement of the same particle in consecutive frames. The intensity map was generated by superimposing the trajectories of beads. Flow velocity and direction maps were computed by averaging the velocities of all particles flowing through the same pixel. All the image reconstruction and processing were performed with custom code in MATLAB (MathWorks, MATLAB R2020b, USA).

### Microfil dye filling

The carotid artery was cannulated with a catheter (Docoll, No. PI-FL140) and Microfil was gradually injected until the whole arterial network was filled with the dye.

### Sensory stimulation

Awake sensory stimulation was carried out by air puffs onto the contralateral vibrissae, where pressure was controlled by a Toohey Spritzer Pressure System IIe (Toohey, Fairfield) and timed triggering by multichannel systems (STG 4002) coupled into the ScanImage software’s frame clock.

Electrical stimulation was performed using subcutaneous needle electrodes (amplitude of 400 µA, 4Hz) inserted into the contralateral whisker pad or hindlimbs and triggered by a Multichannel Systems stimulus generator (STG 4002).

### Collateral occlusion using FeCl_3_

A glass micropipette was filled with a 10% FeCl_3_ solution. The tip of the micropipette was lowered onto the surface of the collateral without piercing the vessel. By capillary action the FeCl_3_ solution caused clot formation inside the collateral after 5-10 minutes.

### Drug applications

BaCl_2_ (30 µM in Ringer solution) was applied topically onto the exposed pia for 10-15 minutes. For imaging the BaCl_2_ (30 µM in Ringer) solution was mixed with 1.5% agarose (SigmaAldrich, Cat.No. A9539) and then applied to the exposed cortex after cooling down to 35°C. 1 mM APV (Tocris, No. 0106) and 250 μM CNQX (Tocris, No. 0190) were injected intracortically (150 nL) using a microinjector and glass micropipettes.

## Acknowledgments

The authors are grateful to Dritan Agalliu for supplying the Cldn5-eGFP mouse line.

## Author contributions

Conceptualization: CG and MEA

Methodology: CG, MW, AVP

Investigation: CG, MW, JD, JC, AVP, QZ, CZ, AVP and MEA

Visualization: CG, ZC, QZ, MW, AVP, NR and MEA

Funding acquisition: BW and MEA

Supervision: BW and MEA

Writing – original draft: CG, MW, AS, SW, BW and MEA

## Data and materials availability

All data are available in the main text or the supplementary materials.

Supplementary Information is available for this paper.

Correspondence and requests for materials should be addressed to Mohamad El Amki.

---

**Extended Data Fig. 1.**
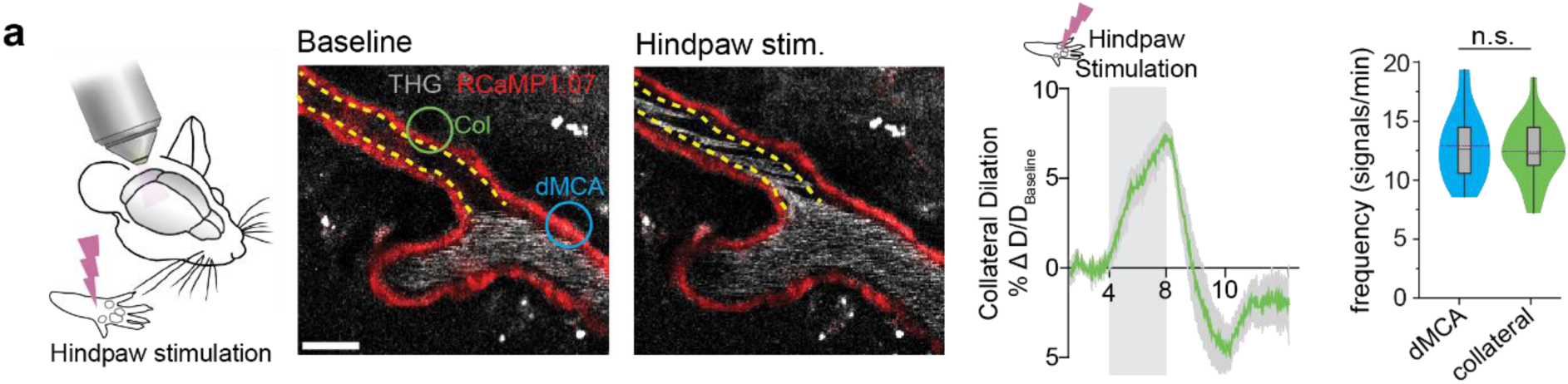
Pial collateral Ca^2+^ - dynamics and involvement in neurovascular coupling. **(a)** Two-photon images of a MCA-collateral branch during hindpaw stimulation. Yellow dotted lines indicate the outline of the vessel segments and circles indicate position of ROIs for Ca^2+^ imaging. On the right, quantification of the collateral dilation during hindpaw stimulation and baseline Ca^2+^ dynamics comparing distal MCA with collaterals N = 3 mice, 9 vessel segments.

**Extended Data Fig. 2.**
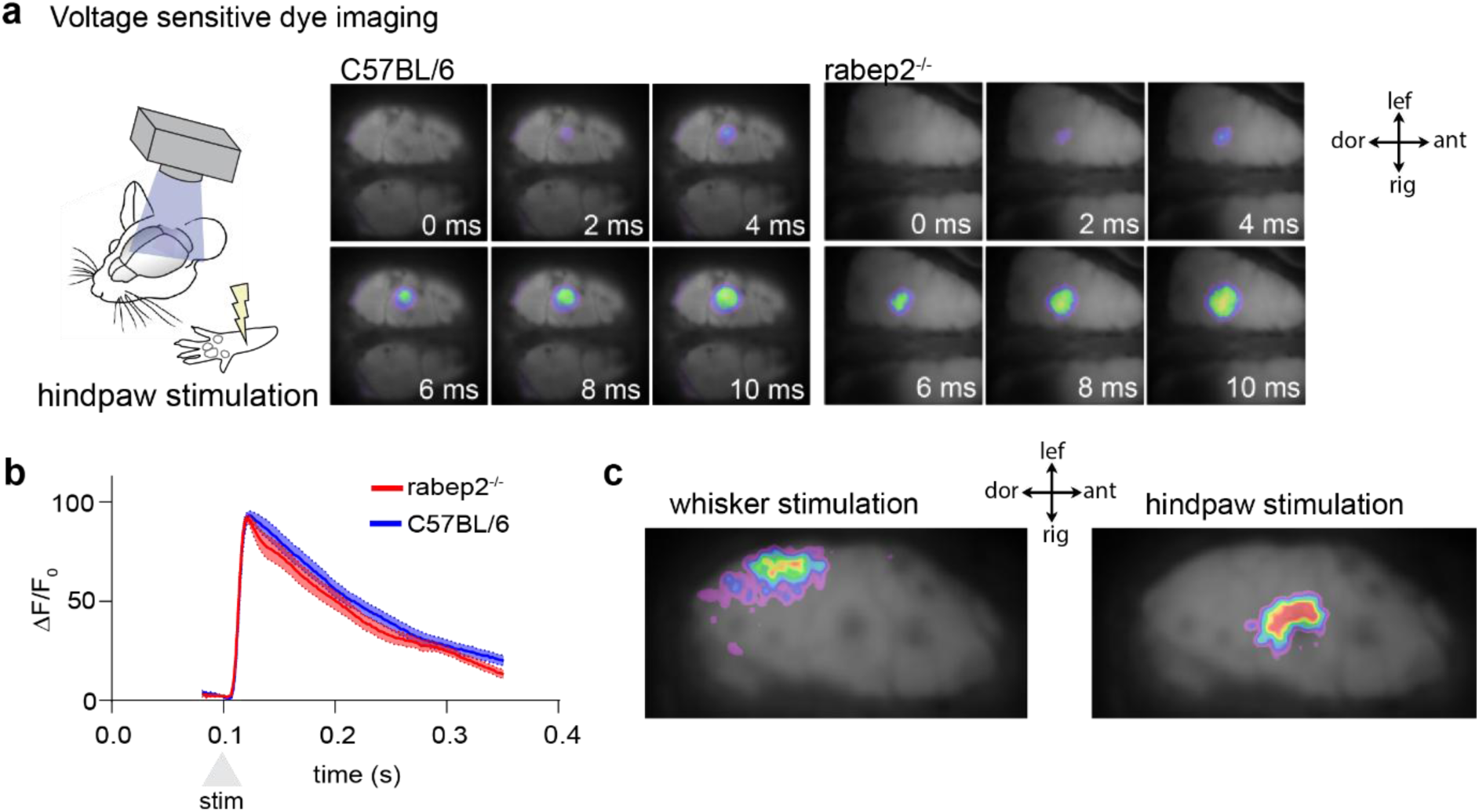
Voltage sensitive dye imaging in C57BL/6 and rabep2^-/-^ mice. **(a)** Timeseries images of VSD recording upon hindpaw stimulation in C57BL/6 and rabep2^-/-^mice. **(b)** Quantification of dF/F_0_ comparing C57BL/6 and rabep2^-/-^ mice. N= 10-11 per group. **(c)** Representative VSD images showing initiation area of whisker and hindpaw stimulation.

**Extended Data Fig. 3.**
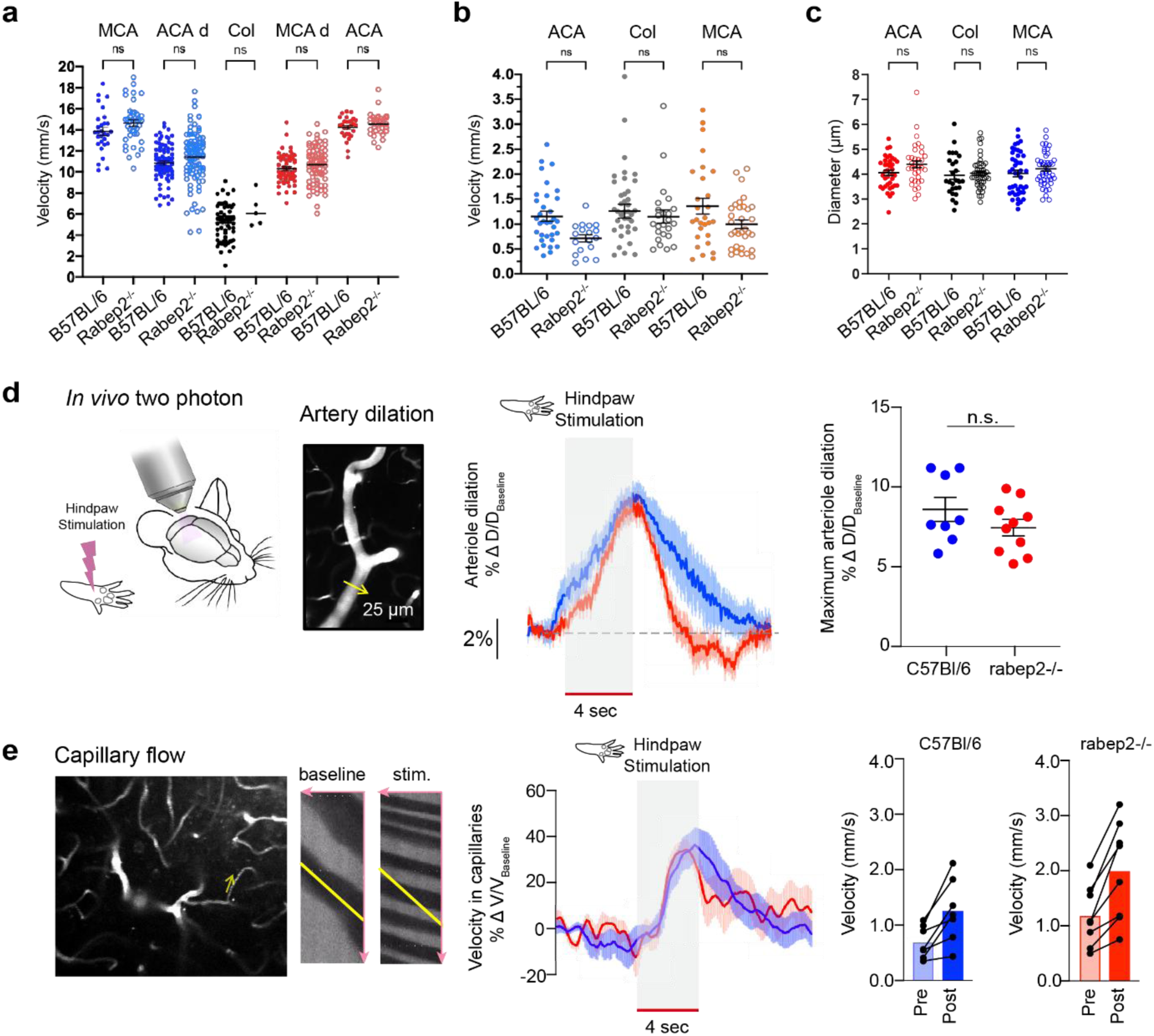
Comparison between C57BL/6 and rabep2^-/-^ blood flow parameters. **(a)** Quantification of PiaFLOW (N = 6-7 mice/group) and 2PM measured blood flow velocities in different vascular branches in awake C57BL/6 and rabep2^-/-^ mice (N = 3-4 mice/group); ns = not significant, d = distal. **(b)** Quantification of 2PM measured blood flow from capillaries connected to MCA, ACA and collateral branches in C57BL/6 and rabep2^-/-^(N = 3 mice/group). **(c)** Quantification of 2PM measured diameters of capillaries connected to MCA, ACA and collateral branches in C57BL/6 and rabep2^-/-^ (N = 3 mice/group). **(d)** Quantification of 2PM measured arteriole dilation in response to hindpaw stimulation in C57BL/6 and rabep2^-/-^ (N = 6-7 mice/group). **(e)** Quantification of 2PM measured blood flow velocities of capillaries connected to upstream ACA arteriole segment in response to hindpaw stimulation in C57BL/6 and rabep2^-/-^ (N = 3 mice/group) mice.

